# A general computational design strategy for stabilizing viral class I fusion proteins

**DOI:** 10.1101/2023.03.16.532924

**Authors:** Karen J. Gonzalez, Jiachen Huang, Miria F. Criado, Avik Banerjee, Stephen Tompkins, Jarrod J. Mousa, Eva-Maria Strauch

**Affiliations:** Institute of Bioinformatics, Franklin College of Arts and Sciences, University of Georgia; Athens, GA 30602, USA; Department of Infectious Diseases, College of Veterinary Medicine, University of Georgia; Athens, GA 30602, USA; Center for Vaccines and Immunology, College of Veterinary Medicine, University of Georgia; Athens, GA 30602, USA; Department of Pathobiology, Auburn University; Auburn, AL 36849, USA; Department of Biochemistry and Molecular Biology, Franklin College of Arts and Sciences, University of Georgia; Athens, GA 30602, USA; Department of Pharmaceutical and Biomedical Sciences, College of Pharmacy, University of Georgia; Athens, GA 30602, USA

## Abstract

Many pathogenic viruses, including influenza virus, Ebola virus, coronaviruses, and Pneumoviruses, rely on class I fusion proteins to fuse viral and cellular membranes. To drive the fusion process, class I fusion proteins undergo an irreversible conformational change from a metastable prefusion state to an energetically more favorable and stable postfusion state. An increasing amount of evidence exists highlighting that antibodies targeting the prefusion conformation are the most potent. However, many mutations have to be evaluated before identifying prefusion-stabilizing substitutions. We therefore established a computational design protocol that stabilizes the prefusion state while destabilizing the postfusion conformation. As a proof of concept, we applied this principle to the fusion protein of the RSV, hMPV, and SARS-CoV-2 viruses. For each protein, we tested less than a handful of designs to identify stable versions. Solved structures of designed proteins from the three different viruses evidenced the atomic accuracy of our approach. Furthermore, the immunological response of the RSV F design compared to a current clinical candidate in a mouse model. While the parallel design of two conformations allows identifying and selectively modifying energetically less optimized positions for one conformation, our protocol also reveals diverse molecular strategies for stabilization. We recaptured many approaches previously introduced manually for the stabilization of viral surface proteins, such as cavity-filling, optimization of polar interactions, as well as postfusion-disruptive strategies. Using our approach, it is possible to focus on the most impacting mutations and potentially preserve the immunogen as closely as possible to its native version. The latter is important as sequence re-design can cause perturbations to B and T cell epitopes. Given the clinical significance of viruses using class I fusion proteins, our algorithm can substantially contribute to vaccine development by reducing the time and resources needed to optimize these immunogens.

## Main

Life-threatening viruses such as the human immunodeficiency virus (HIV)^1^, Ebola virus^2^, Pneumoviruses^3^, and the pandemic influenza^4^ and coronaviruses^5^, use class I fusion proteins to induce the fusion of viral and cellular membranes and infect the host cell. During membrane fusion, class I fusion proteins refold from their metastable conformation (prefusion state) to the highly stable postfusion conformation, likely to provide the energy mediating the fusion reaction^6^. Their essential role in the viral entry as well as their location on the viral surface makes class I fusion proteins one of the major targets of neutralizing antibodies and, thereby, a critical immunogen for vaccination^7^. However, while both pre- and postfusion states are usually immunogenic, the labile prefusion state has been demonstrated to induce a more potent immune response in multiple viral families ^8–12^. Consequently, the prefusion state has become an attractive vaccine candidate when its conformation can be maintained^8,13,14^. Based on structural analyses of the fusion mechanism, the stabilization of the prefusion conformation has been mainly achieved by preventing the release of the fusion peptide or by disrupting the formation of the coiled-coil structure characteristic of the postfusion state^15^. Two strategies have been particularly successful by either designing disulfide bonds at regions undergoing remarkable refolding or introducing proline substitutions to impair the formation of the central postfusion helices^13,16,17^. Other stabilization methods have focused on identifying substitutions that increase favorable interactions or rigidify flexible areas in the prefusion structure. These methods either design cavity-filling substitutions^13,17,18^, neutralize charge imbalances^13,17,18^, or remove buried charged residues^19^. While the strategies mentioned so far have been effective, the lack of an automated approach has limited the number of amino acid changes to be assessed and required extensive testing of variants. Notably, more than one hundred different protein variants were evaluated before finding a stable prefusion conformation of the Filovirus GP protein^20^, the severe acute respiratory syndrome coronavirus 2 spike protein (SARS-CoV-2 S)^21^, and the F protein from Hendra^22^, Nipah^11^, respiratory syncytial virus (RSV F)^17,18^, human metapneumovirus (hMPV F)^23^, and parainfluenza virus types 1-4^8^. To address these limitations, we developed a general computational approach where the fusion protein’s sequence is optimized for the conformation of interest (here, the prefusion state) while destabilizing the other conformation. Our general strategy assumes that conformational rearrangements in class I fusion proteins can be frozen by introducing mutations that reduce the free energy of the prefusion form but do not benefit or better disrupt the postfusion state. While this “negative design” concept has been introduced before in multistate design (MSD) protocols^24^, our efforts to implement leading algorithms in class I fusion proteins, such as the MPI_MSD^25^, evidenced poor sequence sampling. This is likely due to the extensive sequence-structure search space that needs to be evaluated when modeling both states simultaneously of these large, underpacked proteins. Therefore, we modified the design process by avoiding explicit negative design but using the undesired conformation as a guide to identify suboptimal positions. In a second combinatorial design step, we search for an optimal sequence for the conformation of interest within the subset of substitutions identified to improve the prefusion conformation while disfavoring the postfusion conformation. Using this two-step protocol, we are able to control the substitution rates by focusing on the most impactful changes. We have successfully stabilized the prefusion state of several large proteins, namely the RSV F, the hMPV F, and the SARS-CoV-2 spike proteins, illustrating the general use of the method. Importantly, only 3-4 variations were necessary to evaluate experimentally, saving a tremendous amount of time and resources.

## Results and Discussion

### Energy optimization of the prefusion over the postfusion conformation

Fusion proteins must contain various energetically sub-optimal residues for a given conformation to be able to accommodate other conformations required to complete the fusion process^6^. As a first step, we identified these sub-optimal positions for the prefusion conformation based on the protein energetics or its anticipated dynamics (Fig 1). For the first approach, mainly used for the stabilization of the RSV F protein (based on the A2 strain, as published under the PDB 5w23^26^), we uncovered residue positions with contrasting stability between the pre- and postfusion conformations by calculating the energetic contribution of every residue to each state. *In-silico* alanine mutagenesis allowed us to quickly identify contributions towards Gibbs free energy (ΔΔG) of a given residue side chain, approximating the role of the position on the stability of each state^27^. Negative ΔΔG scores (< -1.0 in Rosetta energy units, REU) indicated structural stabilization while positive ΔΔG scores (>1.0 REU) suggested destabilization^27^. Thereby, we created two energetic maps, one for each state, to spotlight residue positions with differential contributions on each conformation (Fig 1). In all our examples, about 40 - 50 positions displayed higher stability in the prefusion state than in the postfusion state.

**Figure 1.**
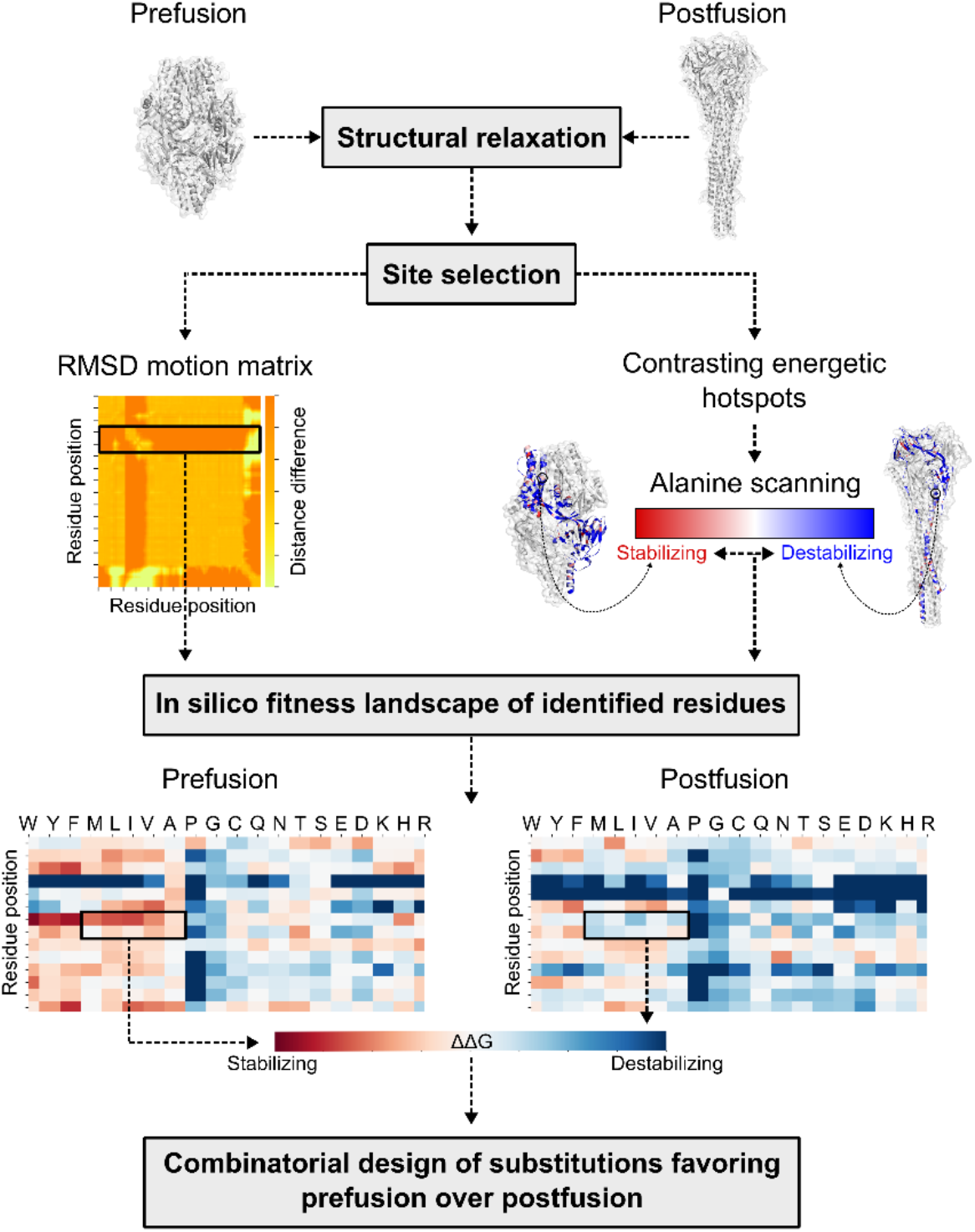
Computational design overview. Step-by-step diagram illustrating the key components of redesigning viral class I fusion proteins. Both pre- and postfusion conformations are input into the pipeline to identify substitutions that favor the prefusion state over the postfusion conformation. RMSD=Root mean square deviation; ΔΔG=change in delta Gibbs free energy.

When alanine scanning was not sufficient to locate meaningful designable spots, as defined by a differential of at least 2 REU, we focused on the dynamics of the protein as a second approach to identify sub-optimal positions. For hMPV F and SARS-CoV-2 S proteins, we defined as “designable” all regions undergoing drastic structural rearrangements between states. Highly movable residues and all positions identified by alanine scanning were exhaustively explored to find substitutions that invert the energetics between states. As the final objective was to find mutations working synergistically rather than individually, all substitutions favoring the prefusion state over the postfusion conformation were assessed in a combinatorial design step (Fig 1). Though several sequences were found to lower the prefusion state while increasing the postfusion state energy, the number of mutations introduced was high (∼40 substitutions). Therefore, to prevent changes in the immunological properties of the proteins, the number of designable positions was decreased. We aimed to introduce less than 10 mutations by focusing on the lowest energy interactions and the preservation of newly introduced hydrogen bonds and salt bridges in the prefusion state, or substitutions with a strong negative effect on the postfusion structure’s energy. The combinatorial design of these substitutions resulted in energetic gaps of at least 119 REU (Fig S1 and Table S1).

### Biochemical characterization of RSV F, hMPV F and SARS-CoV-2 S variants

After ranking all designed sequences based on their energy gap between states, the top 3-4 designed variants were expressed and purified. For RSV F, one (R-1b) out of three designs was found to be a monodispersed and trimeric protein, as evaluated by size exclusion chromatography (SEC) (Fig 2.A). For the hMPV F and SARS-CoV-2 S, two (M-104 and M-305) out of four, and three (Spk-M, Spk-F, and Spk-R) out of three redesigned proteins behaved similarly, respectively (Fig 2.D, G). Remarkably, the hMPV F variant M-104 showed high expression yields, with a 5.5-fold increase with respect to its parent construct the semi-stabilized hMPV F 115-BV^23^ (Fig S2). The structural state of these expressed constructs was then evaluated based on their conservation of prefusion-specific epitopes. As predicted, all designs presented a prefusion-like structure as they tightly bound to prefusion-specific binders such as the RSV F antibodies D25^28,29^ and AM14^29,30^ (Fig 2.B and Fig S3.A), the hMPV F antibodies MPE8^31^ or 465^32^ (Fig 2.E and Fig S3.B), or the angiotensin-converting enzyme 2 (ACE2)^33^ for the SARS-CoV-2 S protein (Fig 2.H).

**Figure 2.**
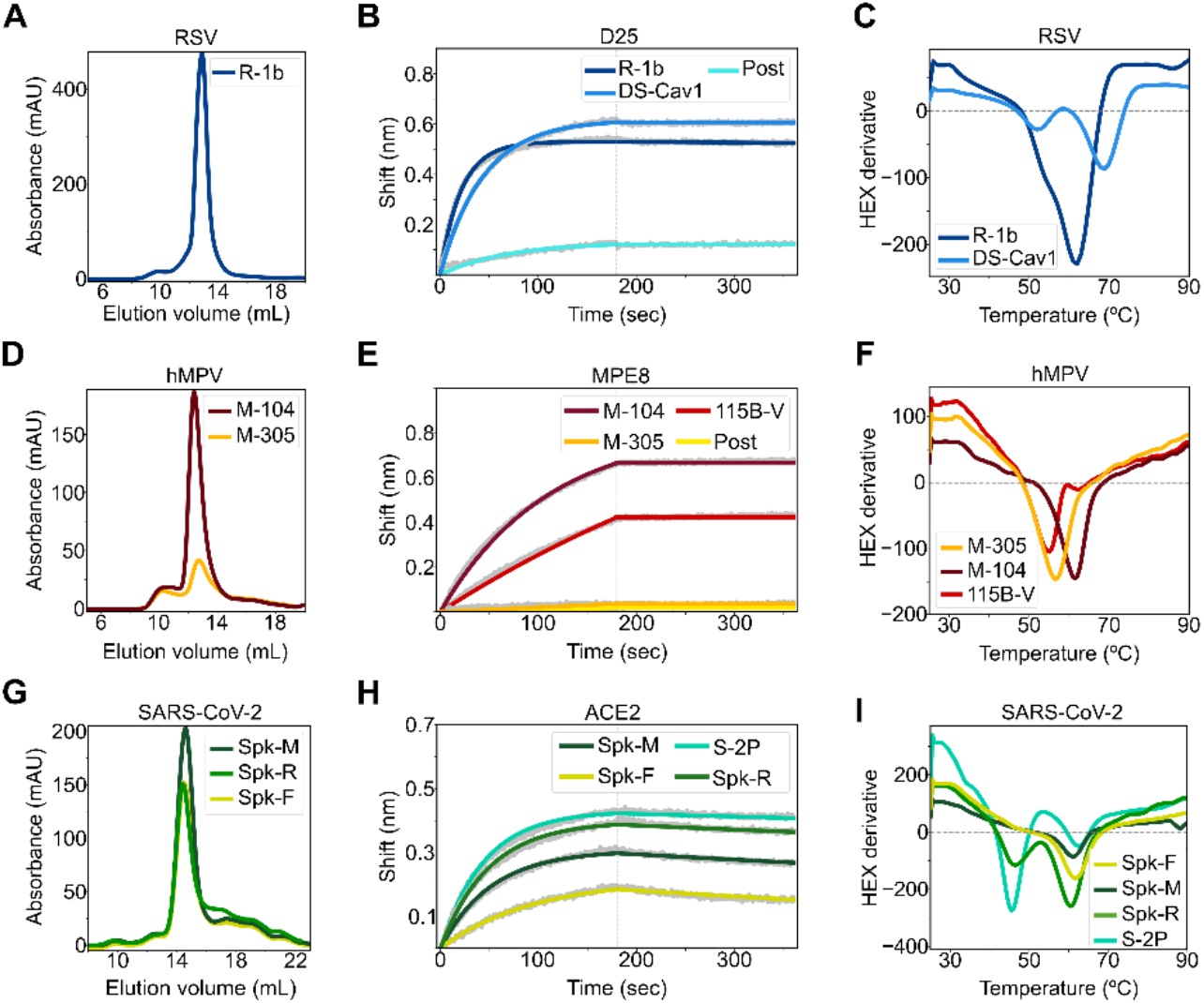
Biochemical characterization of designed variants. **(A)** Size-exclusion chromatography (SEC) of monodispersed RSV F designs. **(B)** Binding of design R-1b to the prefusion-specific antibody D25 compared to the clinical candidate DS-Cav1 and the postfusion RSV A2 F (post). **(C)** Differential scanning fluorimetry (DSF) of design R-1b and the clinical candidate DS-Cav1. DS-Cav1 was used to compare the stability of R-1b as the parental sequence of the latter is not prefusion-stabilized. **(D)** SEC of monodispersed hMPV F designs. **(E)** Binding of designed hMPV F variants to the prefusion-specific antibody MPE8 compared to their parent prefusion construct 115B-V and the postfusion hMPV B2 F (post). **(F)** DSF of designed hMPV F variants and their parent prefusion construct 115B-V. **(G)** SEC of monodispersed SARS-CoV-2 S designs. **(H)** Binding of designed SARS-CoV-2 S variants to ACE2 compared to their parent prefusion construct S-2P. **(I)** DSF of designed SARS-CoV-2 S variants and their parent prefusion construct S-2P. Antibody binding assays show in grey the raw data, in colors the fitted curves, and in dotted lines the end of the association time. Binding constants are shown in Tables S2, S3, and S4.

All expressed proteins showed improved thermal stability compared to their parent prefusion constructs (Table S1). The spike variants Spk-M and Spk-F displayed the highest melting temperature improvement, with ∼15°C increment above the current SARS-CoV-2 vaccine, the S-2P construct^34^ (Fig 2.I). Unlike the vaccine S-2P^34^, the design Spk-M preserved the prefusion conformation even after one hour heating at 55°C as evidenced by ACE2 binding at this temperature (Fig S4.A). This stability seems to compare with the highly stable HexaPro construct, which was achieved by introducing several proline substitutions and experimentally evaluating 100 variants^21^.

A similar scenario was observed in the hMPV F variant M-104; its stability is comparable to variants containing additional disulfide bonds^35^. In fact, M-104 has a higher melting point (61.5°C, Fig 2.F) than the hMPV F variant DS-CavEs (60.7°C)^35^ which has two designed disulfide bonds. Additionally, M-104 remains antigenically unaltered after heating at 55°C (Fig S4.B), as has been seen in DS-CavEs2^35^ which contains four new disulfide bonds. Although, testing under identical conditions is required to validate these comparisons, these results demonstrate how the correct placement of electrostatic interactions can lead to highly stable proteins.

For RSV F, the improvement of the prefusion state stability cannot be well estimated as we started with the wild-type sequence^26^ whose instability substantially impedes its production as an isolated soluble prefusion-state protein^17^; all purified RSV F molecules are found mostly in its postfusion state^36^. Therefore, obtaining the R-1b variant with a melting temperature of 62°C revealed an effective optimization of the sequence to stay in the prefusion conformation (Fig 2.C); especially since no disulfide bonds were introduced and stabilization was achieved through the optimization of non-covalent interactions. Remarkably, design R-1b proved to be antigenically intact even after heating at 55°C (Fig S4.C).

### Structure determination of leading RSV F, hMPV F and SARS-CoV-2 S variants

Negative-stain electron microscopy (EM) confirmed homogeneous trimeric prefusion morphology of leading candidates for all three different fusion proteins studied (Fig S5). We therefore proceeded to obtain atomic details by x-ray crystallization and cryo-EM. The crystal structure of the variants R-1b and M-104 confirmed their prefusion conformation at a resolution of 3.1Å and 2.4Å, respectively (Fig 3.A-B). The accuracy of our computational predictions was reflected on the high structural similarity between the determined structures and the computational models, with root-mean-square deviations (RMSDs) of only 1.193Å (405 Cα atoms) for R-1b, and 0.53Å (416 Cα atoms) for M-104. The 3D classification performed on the spike cryo-EM images also verified the prefusion structure of the protein with particles displaying one receptor binding domain (RBD) in the up conformation (Fig. S6, Table S6). Solving the structure at 3.7Å resolution, revealed that the S2 subunit, which was the only part engineered, agreed closely with the computational model with a RMSD of only 1.345Å (377 Cα atoms) (Fig 3.C).

**Figure 3.**
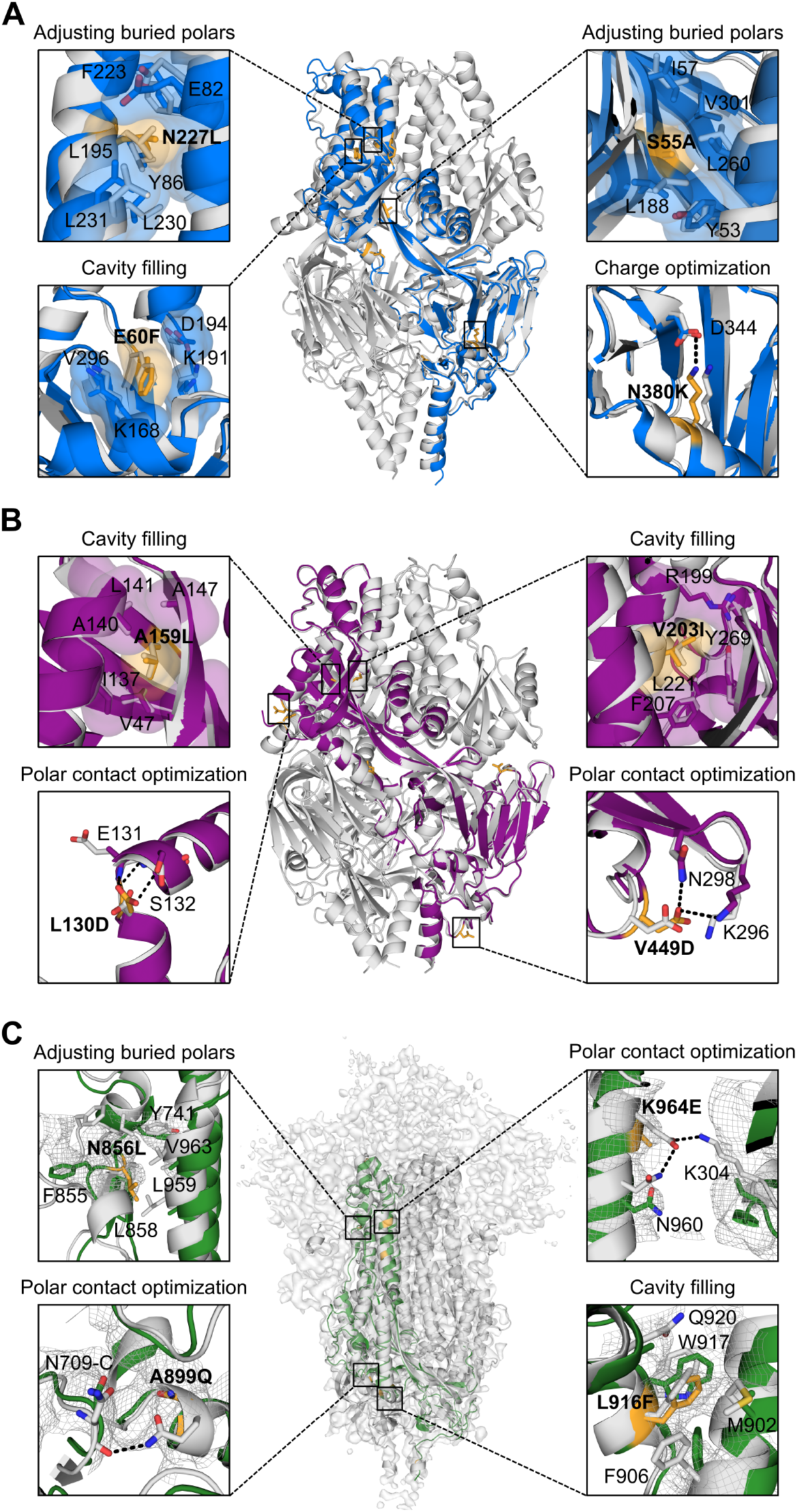
Exemplary stabilizing substitutions of leading designs. **(A)** R-1b. **(B)** M-104. **(C)** S2 subunit of Spk-M with cryo-EM map. The computational model of each protein is displayed as a trimeric structure in grey while the crystal structures and cryo-EM reconstruction model are displayed as monomeric structures in blue (RSV), magenta (hMPV), or green (SARS-CoV-2). Each panel shows a magnified view of selected stabilizing substitutions, featured in yellow sticks, aligned with their computational model. Residues involved in packing changes are displayed with translucent molecular surfaces, and black dotted lines represent hydrogen bonds or salt bridges. As density is missing in the overall map to assign the precise location of the side chains, we displayed existing density as mesh representation to compare agreement with the computational model as stabilized regions have more density than the remainder of the map. The stabilization mechanism of all designed substitutions is presented in Table S1.

Although no significant perturbations were observed in all our variants overall, subtle differences at the antigenic site Ø were identified between R-1b and its parent RSV F protein. Specifically, the α4 helix in R-1b is bent towards residue D200 when compared to the parent RSV F protein. As the antigenic site Ø is intrinsically flexible, structural variations in this region are expected and have been reported in several prefusion-stabilized RSV F proteins, including the clinical candidate DS-Cav1^13,17,18^. For future stabilization efforts, this could be another area of interest for stabilization.

The crystal structures of R-1b and M-104 revealed that despite some deviations between the designed and crystal structure, most introduced substitutions followed their predicted stabilization mechanism by either filing cavities or increasing intra- or interprotomer hydrogen bonds and salt bridges (Fig S7-8 and Table S1). Especially precise rotamer agreement between our computational models and the experimental data was found in cavity-filling mutations, such as E60F in R-1b, and the A159L and V203I in M-104 (Fig 3.A-B). Significant rotamer agreement was also observed in substitutions strengthening polar interactions, such as the N380K in R-1b, and the L130D, V430Q, and V449D in M-104 (Fig 3.A-B and Fig S8). Other mutations such as the S150E and E487N in R-1b did not interact with the predicted residues but still contributed to the prefusion stability by increasing polar interactions at the protomers’ interface (Fig S7). Lastly, we observed the alleviation of buried polar residues by exchanging them for hydrophobic amino acids and improving the overall packing density. This effect was observed, for instance, after swapping the N227 into a leucine residue or removing the unsatisfied hydroxyl of S55 by replacing it with an alanine residue (Fig 3.A). Similarly, we observed these mutations for the hMPV F protein. Finally, the design Spk-M was stabilized by four substitutions filling cavities and five substitutions increasing polar interactions at the S2 subunit, three of which were interprotomer contacts (Fig 3.C, S9 and Table S1).

Although most of the designed substitutions stabilized the prefusion state, the mutations N175R in R-1b, L130D in M-104, and T941D in Spk-M were intended to disrupt the postfusion conformation (Fig S10). These residues are solvent-accessible in the prefusion state and therefore should not have an impact on the stability of that conformation. However, the postfusion conformation places these residues at the six-helix bundle where unsatisfied polar residues are highly unfavored and thereby likely to disrupt the core. As we had the postfusion-specific antibody 131-2A^37^, we sought to prove this hypothesis and confirm that we had not only stabilized the prefusion state but also indeed destabilized the postfusion state. The diminished binding of the 131-2A antibody to R-1b after heating (60°C) (Fig S4.D), which should have converted the protein into its postfusion state, proofed that we fulfilled the design objectives we set out for.

### Immunogenicity of design R-1b

We selected the R-1b variant for a vaccination study due to the availability of a highly stable prefusion control, such as the clinical candidate DS-Cav1^13^. Therefore, to investigate the effect of the introduced mutations on the RSV F immunogenicity, female BALB/c mice were vaccinated twice with either 0.2 or 2 µg of purified R-1b or DS-Cav1 with or without AddaVax adjuvant. Mice were bled at three and nine weeks post-second immunization (Fig 4.A). Sera analysis for binding to prefusion RSV F and RSV A2 neutralization revealed that R-1b induced similar levels of RSV F-specific antibody titers (Fig 4.B-C and Fig S11) and comparable neutralizing activity related to DS-Cav1 (Fig 4.D).

**Figure 4.**
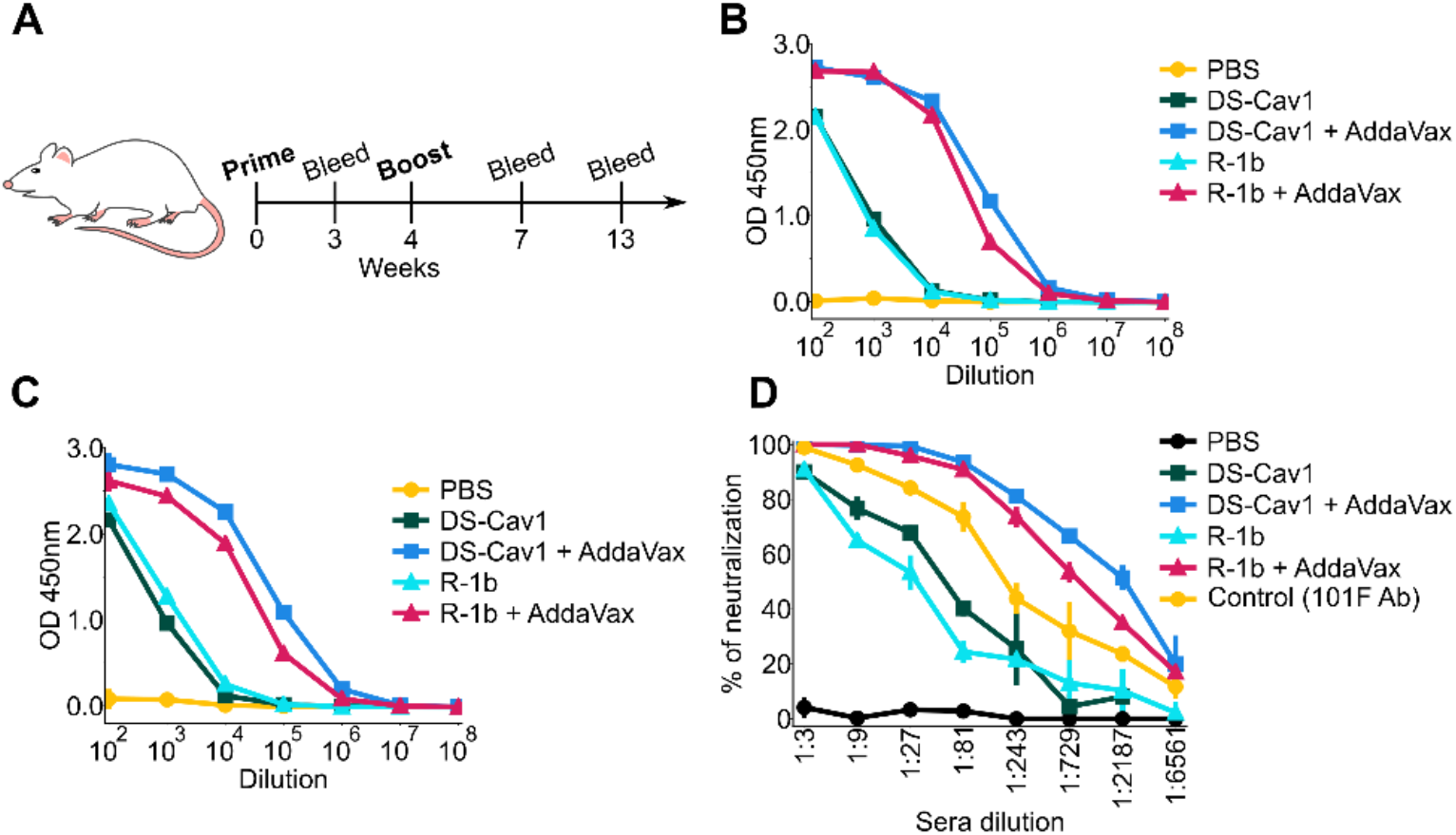
Immunogenicity assessment of R-1b in a mouse model using 0.2ug doses. **(A)** Schematic diagram of vaccination study schedule. **(B)** Serum RSV-specific IgG measured by ELISA three weeks post-boost. **(C)** Serum RSV-specific IgG measured by ELISA nine weeks post-boost. **(D)** Serum neutralization titers determined using RSV A (rA2 strain L19F) and sera from mice nine weeks post-boost. Vertical lines represent the standard deviation of three repetitions using pooled serum samples from mice in each immunization group (5 animals/group).

### General impact

Detailed antibody response studies have illustrated that prefusion-stabilized class I fusion proteins are potent immunogens and promising vaccine candidates. This has been proven to be true for several viruses, including RSV^13,17,18^, hMPV^35^, parainfluenza^8^, Nipah^11^, MERS-CoV^16^, and SARS-CoV-2^21^. Several of these immunogens have been developed by many steps of iterations of manual structure-based design with experimental evaluation, often testing hundreds of combinations of mutations^8,11,17–19,21–23^. To alleviate this laborious exploratory testing, we automated one of the underlying principles behind their stabilization efforts that considers the biophysics of the fusion protein and its large irreversible switch. We developed a computational approach that seeks to freeze the prefusion conformation by learning about suboptimal contacts from its alternate conformation. Our algorithm allows the automated identification of these regions, and their potential substitutions, based on energy differences and relative motion between the two states. We acknowledge that our computational approach can be limited by the need for both pre- and postfusion structures. However, we believe that the high accuracy of current protein structure prediction algorithms could alleviate this drawback^38^. The efficiency of our method has been demonstrated on three different fusion proteins, such as the RSV F, hMPV F, and the SARS-CoV-2 S, where only 3-4 variants were necessary to successfully find a stable prefusion design. Additionally, we were able to validate the immunogenicity of one design in a mouse model, showing similar in vitro neutralization and specific serum IgG patterns compared to a clinical candidate. Therefore, our algorithm could highly impact the vaccine development field by allowing a rapid optimization of both novel class I fusion proteins as well as leading vaccine immunogens.

## Materials and Methods

### I. Computational approach to stabilize the prefusion state

All computational analysis were performed with the Rosetta version: 2020.10.post.dev+12.master.c7b9c3e c7b9c3e4aeb1febab211d63da2914b119622e69b

#### Structure preparation

##### RSV F

All computational analyses were performed on the crystal structure of the RSV F protein in the prefusion (PDB:5w23)^26^ and postfusion (PDB: 3rrt)^36^ conformations. To remove small clashes in the structures, both conformations were refined with the Rosetta relax application guided by electron density data^39^. Density maps were generated from the corresponding map coefficients files associated to the PDB accession codes. These coefficients were transformed into density maps using the Phenix software version 1.15^40^ and the option “create map from map coefficients” (region padding= 0, and grid resolution factor= 0.3333). To include the electron density in the refinement process, the density energy term was activated in the Rosetta scoring function with a weight of 20. This weight was selected given the low-resolution of the density map and the starting structures. During the relaxation protocol, four rounds of rotamer packing and minimization were performed with gradual increases to the repulsive weight in the scoring function^41^. After 5 cycles of relaxation, the quality of the resulting models was evaluated with the Molprobity web service^42^. The structures with the lowest Rosetta energies and Molprobity scores were used for mutational analysis.

##### hMPV F

As described in the RSV F example, the hMPV F prefusion (PDB: 5wb0)^23^ and postfusion (PDB: 5l1x)^43^ conformations were relaxed using their respective electron density data. Due to the high resolution of the starting prefusion hMPV F structure (2.6Å), the weight of the density energy term was increased to 50 to encourage a good agreement with the density map. All other parameters and postprocessing followed the RSV F example.

##### SARS-CoV-2 S

The input SARS-CoV-2 S prefusion and postfusion structures and their corresponding cryo-electron microscopy (cryo-EM) maps were retrieved with the PDB accession codes 6vxx^44^ and 6xra^45^, respectively. Since both structures were not completely solved, missing regions were modeled using the default comparative modeling protocol available in Rosetta^46^. However, the cryo-EM density maps of the input pre- and postfusion structures were integrated in the modeling process to avoid large deviations from the original configuration. Templates selected for prefusion modeling corresponded to the PDB IDs 6m0j^47^ and the initial 6vxx, while the postfusion structure was modeled using 6lxt^48^ and the initial 6xra. Model selection was based on overall agreement with the starting structure and templates, as well as a low Rosetta energy score. These homology models were then relaxed with the RosettaScripts^49^ framework, incorporating fit-to-density parameters established for cryo-EM density^39^. Due to the high resolution of the starting structures, the refinement process was performed with a density scoring weight of 50 and three cycles of FastRelax^41^ in cartesian space. The structures with the lowest Rosetta energies and Molprobity scores were used for mutational analysis.

#### Selection of target positions to redesign

Residue positions to redesign were selected based on two independent approaches: I) contrasting energetic contributions to the pre- and postfusion conformations, and II) location on regions displaying drastic rearrangements between the pre- and postfusion states.

##### Selection based on energetic contributions

Amino acid positions with contrasting energetic contributions between conformations offer an opportunity to manipulate the energetics of the conformational switch by allowing the optimization of one state while the other state is disfavored. Identification of these target spots was done by *in-silico* alanine mutagenesis where the energetic role of each residue on each conformation was estimated using the change in folding energy upon mutation. Consequently, residue positions displaying simultaneous stabilization of the prefusion state and destabilization of the postfusion state were selected as hotspots to redesign. Details about the selection process are described at the computational alanine scanning section

##### Selection based on protein dynamics

When alanine scanning was not enough to identify positions with contrasting stability between conformations, all regions involved in the refolding process were chosen as targets to redesign. To identify these flexible areas, the root mean square deviation (RMSD) of each cα atom was calculated using their corresponding position in the pre- and postfusion structures. Both structures were structurally aligned prior to the analysis, and residue positions displaying motion levels of at least 10Å were selected to redesign. Furthermore, residues flanking the highly movable areas were also considered to redesign when their secondary structure differed between the pre- and postfusion states. Flanking residues were included until a set of 8 (SARS-CoV-2 S) or 16 (hMPV F) consecutive residues matched their secondary structure in the pre- and postfusion structures.

#### Computational alanine scanning

A computational alanine scanning was performed on the pre- and postfusion states of RSV F, hMPV F, and SARS-CoV-2 S proteins to determine the energetic contributions of each amino acid to each conformation. Since the prefusion SARS-CoV-2 S contains domains that are not present in the postfusion conformation, alanine scanning in this protein was limited to shared regions between states. Using the Rosetta ΔΔG protocol “cartesian_ddg”^27^, the backbone and sidechains around the position to be mutated were optimized in the cartesian space and the change in folding energy (ΔG) was computed before and after each alanine substitution^27^. The contribution of every residue to stability was calculated in terms of ΔΔG scores (ΔG _mutant_ Δ-G _wild type_) where alanine changes holding ΔΔG scores < -1.0 were considered stabilizing substitutions while ΔΔG scores > 1.0 were considered destabilizing changes^27^. All calculations were repeated 3 times and the average value among repetitions was used as the final ΔΔG score. To enhance the prefusion stability over the postfusion, regions presenting a stabilizing score in the prefusion conformation but not in postfusion conformation were chosen to redesign. Specifically, designable positions were selected based on a) stabilizing ΔΔG score in the prefusion conformation and destabilizing ΔΔG score in the postfusion conformation, b) stabilizing ΔΔG score in the prefusion conformation and neutral ΔΔG score in the postfusion conformation, or c) destabilizing ΔΔG score in the postfusion conformation and neutral ΔΔG score in the prefusion conformation. To restrict the design process to the most relevant spots, positions meeting any of the above criteria were filtered based on an energetic difference of at least 0.7 (ΔΔG _postfusion_ - ΔΔG _prefusion_). Finally, since positions with native alanine are overlooked with this approach, all alanine-bearing spots were also included as targets to redesign.

#### Computational protein design

To determine the amino acid identities likely to invert the energetics of the pre- and postfusion states, target positions to redesign underwent complete *in-silico* saturation mutagenesis as described at the computational alanine scanning section. Subsequently, substitutions favoring the prefusion state over the postfusion state were chosen for combinatorial design through Rosetta modeling. To bias the design process towards mutations displaying preference for the prefusion state with a high energetic difference between states, the weight of each substitution was adjusted in the Rosetta energy function according to a fitness score. Our fitness score compiled the stabilization effect of one mutation in both pre- and postfusion states by subtracting the ΔΔG prefusion score from the ΔΔG postfusion score (ΔΔG _postfusion_ - ΔΔG _prefusion_). Mutations favoring the prefusion state over the postfusion conformation were then characterized by positive fitness scores where higher values represented bigger energetic gaps between states. The fitness score was incorporated into the Rosetta score function through a residue-type constraint term derived from the “FavorSequenceProfile” mover. Since this mover was initially created to re-weight amino acid substitutions depending on their occurrence in a multiple sequence alignment, we have replaced the original position-specific substitution matrix (PSSM) input by a fitness score matrix. To follow the PSSM format, negative fitness scores were replaced by zero and a 0.05 pseudocount was used for the log-odds scores calculation. After tuning the profile weights of each residue, “allowed” mutations at every amino acid position were defined as those with a fitness score greater than or equal to 0.7. For challenging targets such as the SARS-CoV-2 S, the threshold difference was increased to 2 to focus on the most significant substitutions. Likewise, beneficial mutations for both states were allowed only if the stabilization effect in the prefusion state was at least 4 units greater than in the postfusion state. Finally, the combinatorial sequence design was carried out by the FastDesign algorithm^50,51^. Upon conclusion, further optimization of a specific target spot was optionally done by applying FastDesign (all amino acids allowed) on residues neighboring 6Å around the point of interest and limiting packing and minimization to a 12Å sphere. The design process was initially performed on the prefusion conformation, and the resulting sequences were modeled on the postfusion structure for energetic comparisons.

#### Selection of top designs

Promising designs were first sorted based on their Rosetta total energy score. Before comparison, the parent pre and postfusion structures were relaxed and energetically minimized using the same protocol as the designed models. Top candidates corresponded to designs showing a lower energy score in the prefusion state compared to the parent pre- and postfusion conformations. Analogously, the designed sequence in the postfusion state had to display a higher energy score than the parent postfusion conformation. The selected designs were then analyzed at the residue level to identify which mutations contributed more to the energetic gap between states. In this regard, mutations were filtered according to different structural metrics, such as Rosetta per-residue energy, and the number of hydrogen bonds and van der Waals contacts. To improve the prefusion state over the postfusion conformation, mutations showing favorable metrics on the prefusion state and unfavorable values for the postfusion conformation were selected to be tested experimentally.

### II. Protein expression and characterization

#### Protein expression

The top 3-4 RSV F, hMPV F, and SARS-CoV-2 S computational designs as well as the starting constructs hMPV F 115-BV^23^, and SARS-CoV-2 S-2P^34^, and the control variants RSV F DS-Cav1^13^, postfusion RSV A2 F^52^, and postfusion hMPV B2 F^53^ were expressed by transient transfection of FreeStyle 293-F cells (Thermo Fisher) with polyethylenimine (PEI) (Polysciences). All computationally designed variants were produced in pCAGGS plasmids encoding the sequence of interest, a C-terminal T4 fibritin trimerization motif (Foldon), and a His6-tag. The designed RSV F constructs contained residues 1-105 and 137-513, and a flexible linker replacing the furin cleavage sites and p27 peptide (“QARGSGSGR”)^17^. Likewise, the designed hMPV F sequences included residues 1-95 and 103-472, a modified cleavage site “ENPRRRR”, and the A185P mutation^23^. Finally, the designed SARS-CoV-2 S variants followed the semi-stabilized SARS-CoV-2 S-2P protein sequence^34^, with two proline substitutions at residues 986 and 987, and a “GSAS” linker replacing the furin cleavage site. All DNA sequences were codon optimized for human expression using the online tool GenSmart Codon Optimization^54^. Cells were cultured at 37°C and 8% CO_2_, and the culture supernatant was harvested on the 3rd day after transfection. Proteins were purified by nickel affinity chromatography followed by size-exclusion chromatography (SEC) in phosphate-buffered saline (PBS) buffer pH 7.4. RSV F and hMPV F variants were SEC purified using a Superdex200 column (Cytiva) while SARS-CoV-2 S was purified with a Superdex6 column (Cytiva). The angiotensin-converting enzyme 2 (ACE2) was expressed as an Fc-fusion^33^ after transient transfection as described above. The protein was purified using a Protein A agarose gravity column (Millipore Sigma) followed by SEC using a S200 column.

#### Antigenic characterization

Bio-layer interferometry (BLI) was used to evaluate the structural and antigenic conservation of prefusion-specific epitopes. The prefusion-specific binders used for this purpose were the antibodies D25^28,29^ (Thermo Fisher) and AM14^29,30^ (Cambridge Biologics) for RSV F, MPE8^31^ and 465^32^ for hMPV, and the angiotensin-converting enzyme 2 (ACE2)^33^ for SARS-CoV-2 S. All binders were immobilized on Protein A sensors (GatorBio) at a concentration of 15 nM (RSV F, and hMPV F), or 40 nM (SARS-CoV-2 S). Binding against expressed designs was tested at eight different protein concentrations starting from 200 nM (RSV F, and hMPV F) or 400 nM (SARS-CoV-2 S) and decreasing by 1:2 dilutions. All solutions had a final volume of 200 µL/well using PBS buffer supplemented with 0.02% tween-20 and 0.1mg/mL bovine serum albumin (BLI buffer). Biosensor tips were equilibrated for 20 min in BLI buffer before loading of binders. Loading was then carried out for 180s followed by a baseline correction of 120s. Subsequently, association and dissociation between the binders and designed variants were allowed for 180s each. To validate the BLI results, binding with previous prefusion-stabilized proteins such as the RSV F DS-Cav1^13^, hMPV F 115-BV^23^, and SARS-CoV-2 S-2P^34^ was used as positive controls, and binding with the postfusion constructs RSV A2 F^52^, and hMPV B2 F^53^ was used as negative controls. All assays were performed using a GatorPrime biolayer interferometry instrument (GatorBio) at a temperature of 30°C and frequency of 10 Hz. Data analysis was completed with the GatorOne software 1.7.28, using a global association model 1:1.

#### Thermal stability

The thermal stability of the expressed variants was assessed by differential scanning fluorimetry (DSF). The samples were prepared by creating a solution containing 1.2µL SYPRO orange fluorescent dye (Thermo Fisher) with 3 µL of 100mM MgCl_2_, 3 µL of 1M KCl, and 3 µL of 1M Tris (pH 7.4). The final solution volume was 60 µL with a protein concentration of 3.5 µM. A negative sample with no protein was also prepared as background control. All measurements were performed by triplicates using 20 µL of sample. The data was collected with a qPCR instrument (CFX Connect, BioRad) and a temperature ramp from 25 to 90°C with 0.5°C increments. The melting temperature was determined based on the lowest point of the negative first derivative of the SYPRO Orange signal.

Antigenic preservation in variants displaying the highest melting temperature was further evaluated after one hour incubation at 55 and 60°C. This process was carried out in a thermocycler with heated lid (T100, BioRad). The conservation of the antigenic sites was determined by binding to prefusion-specific binders as described at the antigenic characterization section. Conversion to the postfusion state was also evaluated for the RSV F variant R-1b using the postfusion-specific antibody 131-2A^37^ (Millipore Sigma) and the postfusion RSV A2 F^52^ as positive control.

#### Negative-stain electron microscopy

Purified R-1b, M-104, and Spk-M (buffer-exchanged into 50 mM Tris pH 7.5 and 100 mM NaCl) were applied on a carbon-coated copper grids (400 mesh, Electron Microscopy Sciences) using 5 μL of protein solution (10 μg/mL) for 3 min. The grid was washed in water twice and then stained with 0.75% uranyl formate (R-1b) or Nano-W (Nanoprobes) (M-104, and Spk-M) for 1 min. Negative-stain electron micrographs were acquired using a JEOL JEM1011 transmission electron microscope equipped with a high-contrast 2K-by-2K AMT mid-mount digital camera.

#### X-rays crystallization

The trimeric R-1b and M-104 proteins were concentrated to 14 mg/mL and 13.9 mg/mL, respectively, and crystallization trials were prepared on a TTP LabTech Mosquito Robot in sitting-drop MRC-2 plates (Hampton Research) using several commercially available crystallization screens. R-1b crystals were obtained in the Index HT (Hampton Research) in condition H6 (0.2 M Sodium formate, 20% w/v PEG 3,350), while M-104 crystals were obtained in the Crystal screen (Hampton research) in condition C10 (0.1 M Sodium acetate trihydrate pH 4.6, 2.0 M Sodium formate). Crystals were harvested and cryo-protected with 30% glycerol in the mother liquor before being flash frozen in liquid nitrogen. X-ray diffraction data were collected at the Advanced Photon Source SER-CAT beamLine 21-ID-D. Data were indexed and scaled using XDS^55^. A molecular replacement solution was obtained in Phaser^40^ using the prefusion RSV F SC-TM structure (PDB 5c6b)^17^ or the prefusion hMPV F 115-BV (PDB 5wb0)^23^. The crystal structures were completed by manually building in COOT^56^, followed by subsequent rounds of manual rebuilding and refinement in Phenix^40^. The data collection and refinement statistics are shown in Table S5.

#### Cryo-electron microscopy

Spk-M cryo-EM density data was obtained by the Eyring Materials Center at Arizona State University (ASU). Purified protein was diluted to a concentration of 0.35 mg/mL in Tris-buffered saline (TBS) and applied to plasma-cleaned CF-300 2/1 grids before being blotted for 3 seconds in a Vitrobot Mark IV (Thermo Fisher) and plunge frozen into liquid ethane. 3,257 micrographs were collected from a single grid using a FEI Titan Krios (Thermo Fisher) equipped with a K2 summit direct electron detector (Gatan, Pleasantville CA.). Data collection was automated with SerialEM with a defocus range of -0.8 to -2.6 um in counting mode on the camera with a 0.2 second frame rate over 8 seconds and total dose of 58.24 electron per angstrom squared. Images were processed using cryoSPARC V3.3.2^57^ (Fig. S11). Micrographs were patch motion corrected. After particle extractions, the blob picker was used, and 1,551,079 and picking was manually adjusted to reduce “blobs” to 1,394,889 particles. After 2D classifications first 4 classes were ab initio reconstructed and heterogeneously refined. The most populated map was refined with homogenous refinement in cryoSPARC, resulting in a 3.72 Å map. The map was further processed in DeepEM^58^. The resulting final map was aligned with a previously published SARS-CoV-2 S with one RBD domain up (PDB ID 6vyb^44^) using UCSF Chimera-1.15^59^. Mutations and coordinate fitting were done manually using COOT^56^ and structure optimization was achieved by iterative refinement using Phenix real space refinement^40^ and COOT. The model and map statistics are presented in Table S6.

### III. Animal studies (RSV only)

#### Mouse immunization

All animal experiments were performed in accordance with the guidelines and approved protocols by the Institutional Animal Care and Use Committee at University of Georgia, Athens, USA. Six-to-eight-week female BALB/c mice were purchased and housed in microisolator cages in the animal facility at University of Georgia. Food and water were provided ad libitum. After acclimation period, 5 mice per group were intramuscularly (i.m.) inoculated with total 100 µL of two different doses (2 µg and 0.2 µg) of either purified DS-Cav1 or R-1b protein with AddaVax adjuvant (50% v/v) or PBS at weeks 0 and 4 (Prime and Boost Vaccination protocol) (Fig 4.A). Bleeds were collected from tail vein pre- and post-immunization (3, 7, and 13 weeks) and sera were analyzed by ELISA and neutralization assay.

#### Measurement of IgG response by ELISA

Medium binding 96 wells microplates (Greiner Bio-One) were coated with 50 μL per well of DS-Cav1 or R-1b protein at 2 μg/mL at 4°C overnight. Plates were washed in PBS/0.05% Tween 20 (Promega) and then blocked with blocking buffer solution (PBS/0.05% Tween 20 /3% non-fat milk (AmericanBio) /0.5% Bovine Serum Albumin (Sigma)) at room temperature for 2 hours. Pooled serum from each group of mice pre-and post-different stages of immunization or control were inactivated at 56°C per 1 hour for subsequent serial dilution in blocking buffer. 100 µL per well of inactivated diluted sera were incubated in triplicate at room temperature for 2 hours. Subsequently, three washes were performed, and plates were incubated with peroxidase-labeled goat anti-mouse IgG (1:3500) (SeraCare) diluted in blocking buffer. After one-hour incubation at room temperature, plates were washed and TMB substrate working Solution (Vector Laboratories) was added. After 10 min at room temperature, the reaction was stopped by adding 50 μL per well of Stop Solution for TMB ELISA (1N H2SO4). Plates were then read on Cytation7 imaging Reader (BioTek) at 450 nm.

#### RSV neutralization assays

Pooled serum samples from mice in each immunization group (5 animals/group) after vaccination and boost (13 weeks after the beginning of the experiment or prime vaccination/ 9 weeks after Prime and Boost vaccination) were diluted in Opti-MEM media (Thermofisher) in serial 3-fold dilutions. Antibody 101F (provided by Jarrod J. Mousa) was used as a positive control for virus neutralization starting at 20 μg/mL. Further, dilutions were mixed with 120 focus-forming units (FFU) of RSV A virus (strain: rA2 line19F) (kindly provided by Dr. Martin Moore) and incubated for 1 hour at room temperature. Subsequently, RSV and sera/antibody dilutions were added to Vero E6 (ATCC) monolayer (105 cells/well) in triplicate and incubated for one hour at 37°C, gently rocking the plate every 15 minutes. Following the incubation, cell monolayers were covered with an overlay of 0.75% methylcellulose dissolved in Opti-MEM with 2% Fetal Bovine Serum (FBS) (Thermofisher) and incubated at 37 °C, 5% CO2. After four days, the overlay was removed, and wells fixed with neutral buffered formalin 10% (Sigma) at room temperature for 30 minutes. Further, fixed monolayers were washed with water and dried at room temperature. An FFU assay was performed to identify the percentage of RSV neutralization. Briefly, wells were washed gently with PBS-0.05% Tween-20 (Promega) and incubated per one hour with anti-RSV polyclonal antibody (EMD Millipore) diluted 1:500 in dilution buffer [5% Non-fat dry milk (AmericanBio) in PBS-0.05% Tween-20]. Plates were washed three times with PBS-0.05% Tween-20, followed by 30 minutes incubation of secondary antibody HRP conjugate rabbit anti-goat IgG (Millipore Sigma) diluted 1:500 in dilution buffer. After incubation, wells were washed, and TMB Peroxidase substrate (Vector Laboratories) was added for 1 hour at room temperature. The visualized foci per well were counted under an inverted microscope.

## Acknowledgments

We would like to thank Luki Goldschmidt for his support maintaining the computing resources, including CyroSPARC. We would like to thank Dr. Dewight Williams at Arizona State University’s Eyring Materials Center for taking the cryoEM data.

## Author contributions

Conceptualization: KJG, EMS

Methodology: KJG, JH, MFC, AB, EMS

Formal analysis: KJG, JH, MFC, JJM, EMS

Writing–original draft: KJG, EMS

Writing–reviewing & editing: KJG, JH, MFC, SMT, JJM, EMS

Visualization: KJG, EMS

Supervision: SMT, JJM, EMS

Project administration: EMS

Funding acquisition: SMT, JJM, EMS

## Data and materials availability

Atomic coordinates of the reported structures and cryo-EM map were deposited in the Protein Data Bank under accession codes 7TN1, 8E15, and 8FEZ, and in the Electron Microscopy Data Bank under accession code EMD-29035. R-1b, M-104, and Spk-M plasmids are available from EMS under a material transfer agreement with the University of Georgia. Rosetta is available through licensing https://www.rosettacommons.org. Scripts for generating designs will be available on https://github.com/strauchlab/stabilization upon publication.

